# BioNumPy: Fast and easy analysis of biological data with Python

**DOI:** 10.1101/2022.12.21.521373

**Authors:** Knut Rand, Ivar Grytten, Milena Pavlovic, Chakravarthi Kanduri, Geir Kjetil Sandve

## Abstract

Python is a popular and widespread programming language for scientific computing, in large part due to the powerful *array programming* library NumPy, which makes it easy to write clean, vectorized and efficient code for handling large datasets. A challenge with using array programming for biological data is that the data is often non-numeric and variable-length (such as DNA sequences), inhibiting out-of-the-box use of standard array programming techniques. Thus, a tradition in bioinformatics has been to use low-level languages like C and C++ to write efficient code. This makes the tools less transparent to the average computational biologist - making them harder to understand, modify and contribute to.

We here present a new Python package BioNumPy, which adds a layer on top of NumPy in order to enable intuitive array programming on biological datasets. BioNumPy is able to efficiently load biological datasets (e.g. FASTQ-files, BED-files and BAM-files) into NumPy-like data structures, so that NumPy operations like indexing, vectorized functions and reductions can be applied to the data. We show that BioNumPy is considerably faster than vanilla Python and other Python packages for common bioinformatics tasks, and in many cases as fast as tools written in C/C++. BioNumPy thus bridges a long-lasting gap in bioinformatics, allowing the same programming language (Python) to be used across the full spectrum from quick and simple scripts to computationally efficient processing of large-scale data.

## Introduction

Python is one of the most commonly used and fastest growing programming languages [1]. Being a high-level language, Python is flexible and suits a wide variety of analyses. It is both easy to learn for biologists new to programming and a powerful language for experienced bioinformaticians. However, a common hurdle is that vanilla Python is too slow to be a viable option for large-scale analyses. Thus, bioinformaticians often end up using non-transparent and error-prone one-liners on the unix command line, or end up developing and using tools written in low-level languages such as C and C++.

In other scientific fields (e.g. physics, engineering and machine learning), Python is being extensively and successfully used for high-performance computing and large-scale analysis [2, 3, 4, 5]. This is in large part thanks to the very powerful *array programming* package NumPy [6], which enables memory-efficient representation and fast analysis of numeric data (similar to R and MATLAB). For problems that lend themself to the array programming paradigm, solutions based on suited libraries in high-level languages are usually easier to write, read, use and adapt than corresponding programs written in low-level languages like C and C++. However, the discrete and variable-length nature of biological sequence data inhibits the out-of-the-box use of standard array programming languages and libraries. Due to a lack of suited high-performance libraries in Python, two distinct implementation strategies have come to dominate the processing and analysis of biosequence data: a) the use of high-level languages like Python with libraries like BioPython [7] and Biotite [8] for smaller-scale analytical exploration in individual life science investigations, and b) the use of low-level languages like C to create command-line executables for common compute-intensive tasks.

We here present the BioNumPy package, which enables efficient and intuitive array programming on biological data in Python. BioNumPy supports a broad range of bioinformatics analyses, with the main philosophy being that data structures should behave as closely as possible to standard numeric NumPy arrays. This means that BioNumPy is easy to learn for users familiar with NumPy or with array programming languages like R and Matlab. BioNumPy is open source and freely available at https://github.com/bionumpy/bionumpy/, and can be installed through the Python package manager Pip. BioNumPy comes with extensive documentation and a user guide that makes it easy to use for a wide range of bioinformatics problems.

## Results

### The BioNumPy Library

BioNumPy is a Python package for efficiently reading, representing and analysing biological datasets. All time-critical operations are implemented in NumPy, meaning that BioNumPy performs comparably to customised low-level language implementations. The key features of BioNumPy are:

1. Reading/writing biological datasets directly to/from NumPy-like data structures, providing easy access to the data through an intuitive and easy-to-use API.
2. Processing and analysing such biological data efficiently using a NumPy-like interface.

As an example, reading in a set of sequences from a FASTQ-file and computing their GC-content is as simple as:

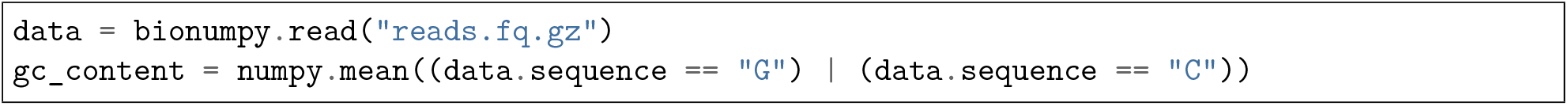

In the example above, the *data*.*sequence* object is a NumPy-like data structure containing all the sequences in our datasets. Common NumPy operations, like indexing and vectorized operations, work with this data structure. BioNumPy also supports broadcasting of functions along one and two-dimensional arrays (in the same way as NumPy), allowing to e.g. easily compute GC content per sequence, or per position across all sequences. This makes BioNumPy powerful and flexible, allowing it to perform a wide range of operations on biological datasets.

BioNumPy comes with extensive documentation, available at https://bionumpy.github.io/bionumpy, showing how to do common tasks for a wide range of data formats and domains.

### Benchmarks

We compare the speed of BioNumPy against other existing Python packages and commonly used non-Python tools on a set of typical bioinformatics tasks. As seen in Figure 1, we find that BioNumPy is generally considerably faster than vanilla Python solutions, including the commonly used Python packages BioPython and Biotite, which mostly rely on Python for-loops to perform operations on datasets. On problems where designated efficient bioinformatics tools are commonly used (intersection of BED-files, kmer counting and VCF operations), we find that BioNumPy is close to, or as efficient as, tools written in C/C++ (BEDTools [9], Jellyfish [10] and BCFTools [11]). While these benchmarks only cover a very small subset of operations, and we only compare against a small subset of available tools, we believe the results still illustrate that BioNumPy can achieve the same performance as dedicated tools written in low-level languages. A Snakemake pipeline for reproducing the results can be found at https://github.com/bionumpy/bionumpy/tree/master/benchmarks, along with an open invitation to expand the benchmark with additional tools and cases.

**Fig. 1:**
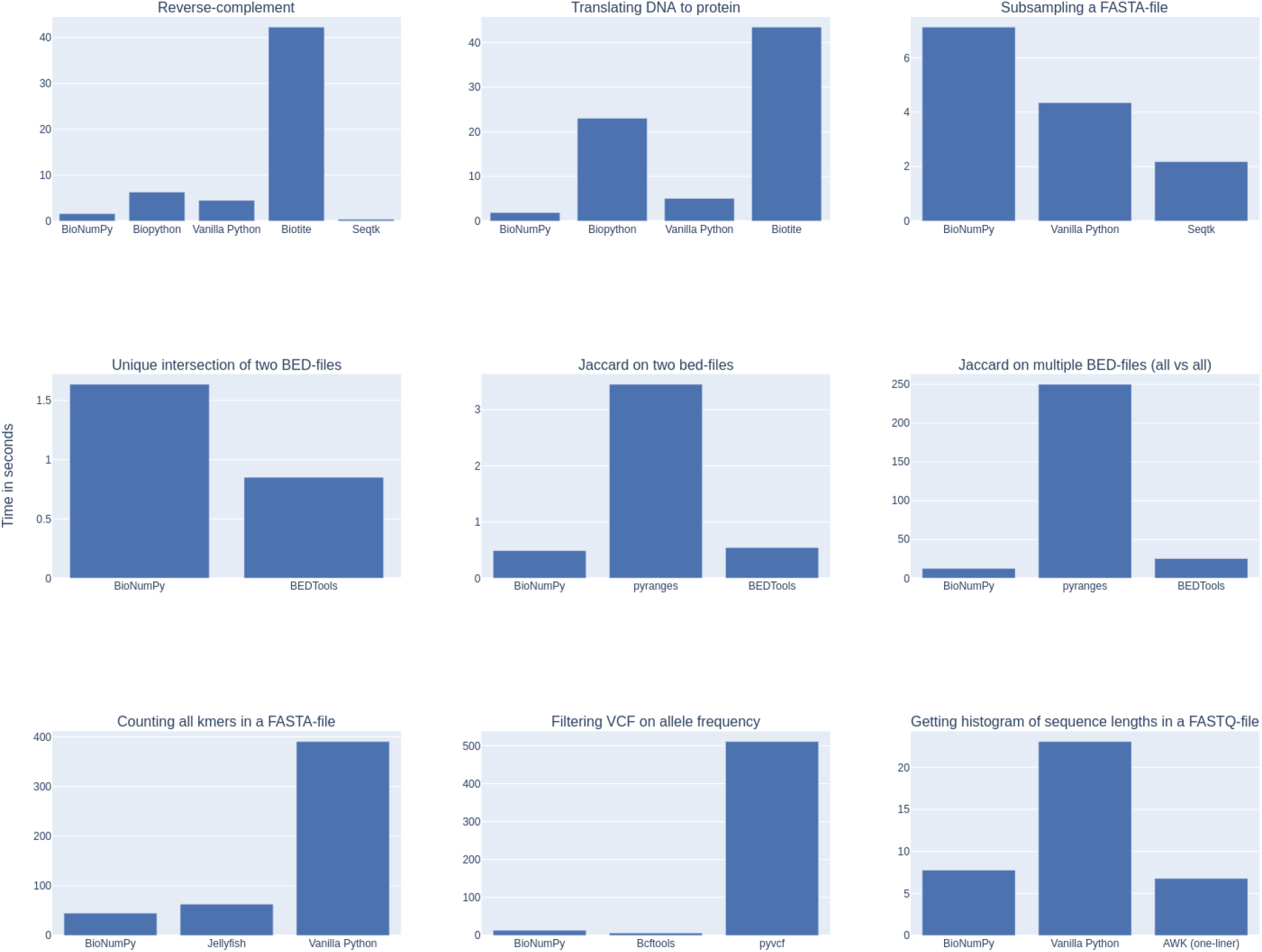
Benchmarking BioNumPy against other tools and methods on various typical bioinformatics tasks.

### Example usage

We here show two examples to illustrate how BioNumPy can be used.

#### Example 1: Using BioNumPy on sequence data

In the following example, we represent a few sequences with BioNumPy and show how basic NumPy functionality works:

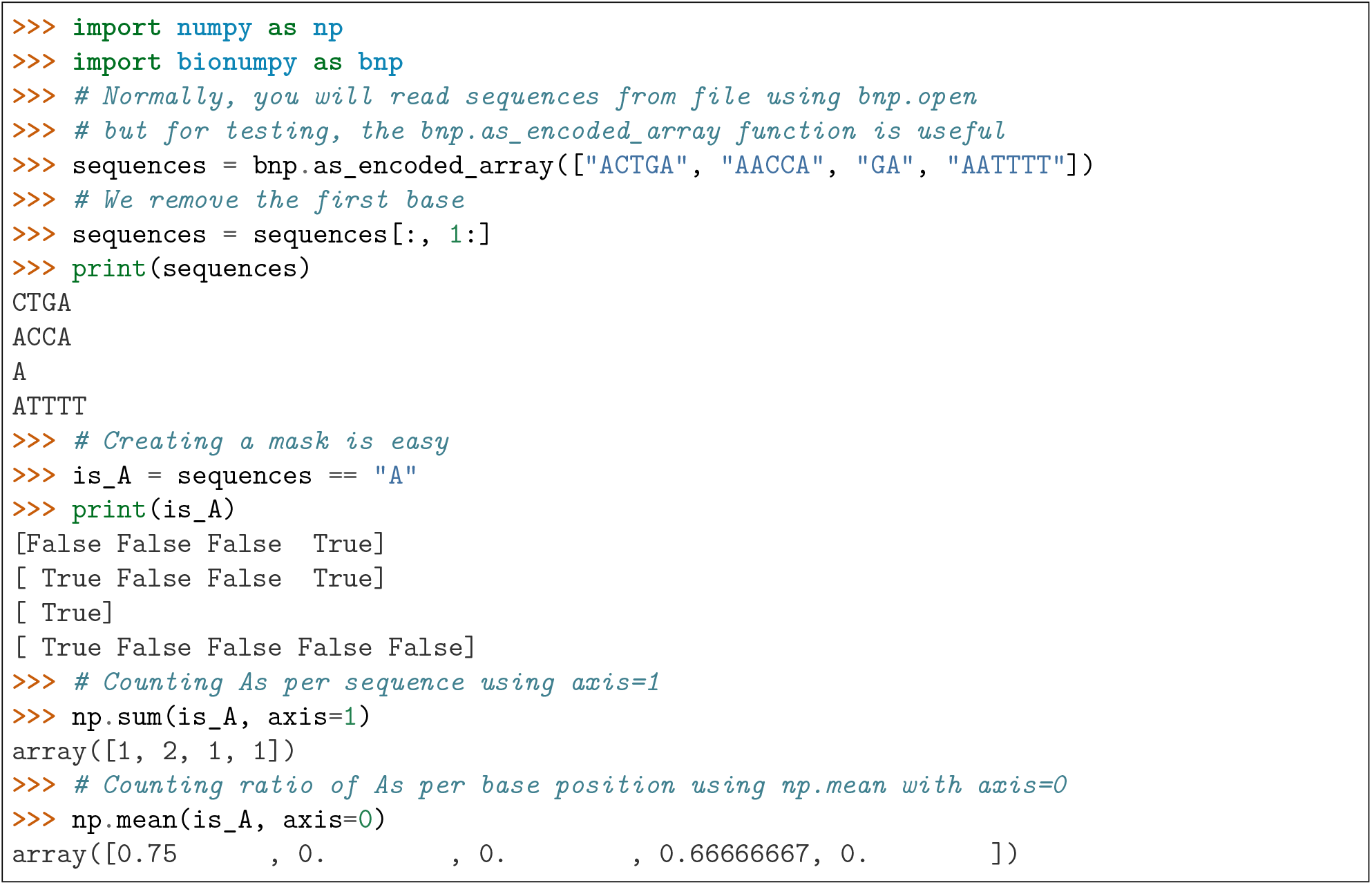

#### Example 2: Analysing motif matches inside transcription factor peaks

In the following example, we show how BioNumPy can be used to easily combine different types of datasets. We read transcription factor peaks from a bed file, fetch the peak sequences from an indexed reference genome and analyse motif scores within the peaks using data from the Jaspar database [12]:

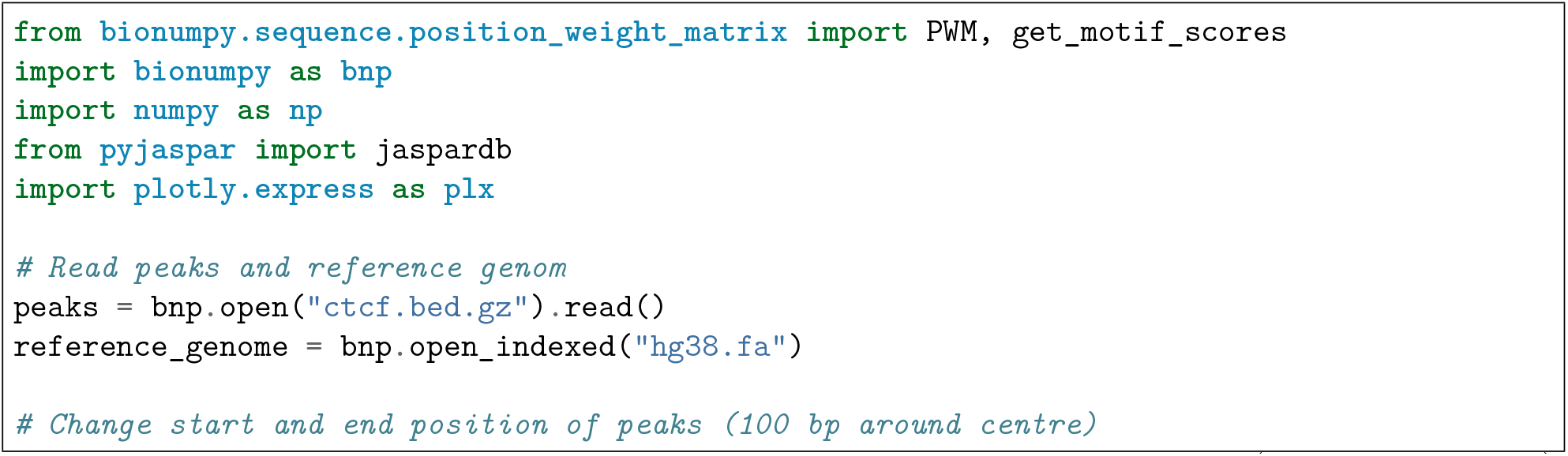
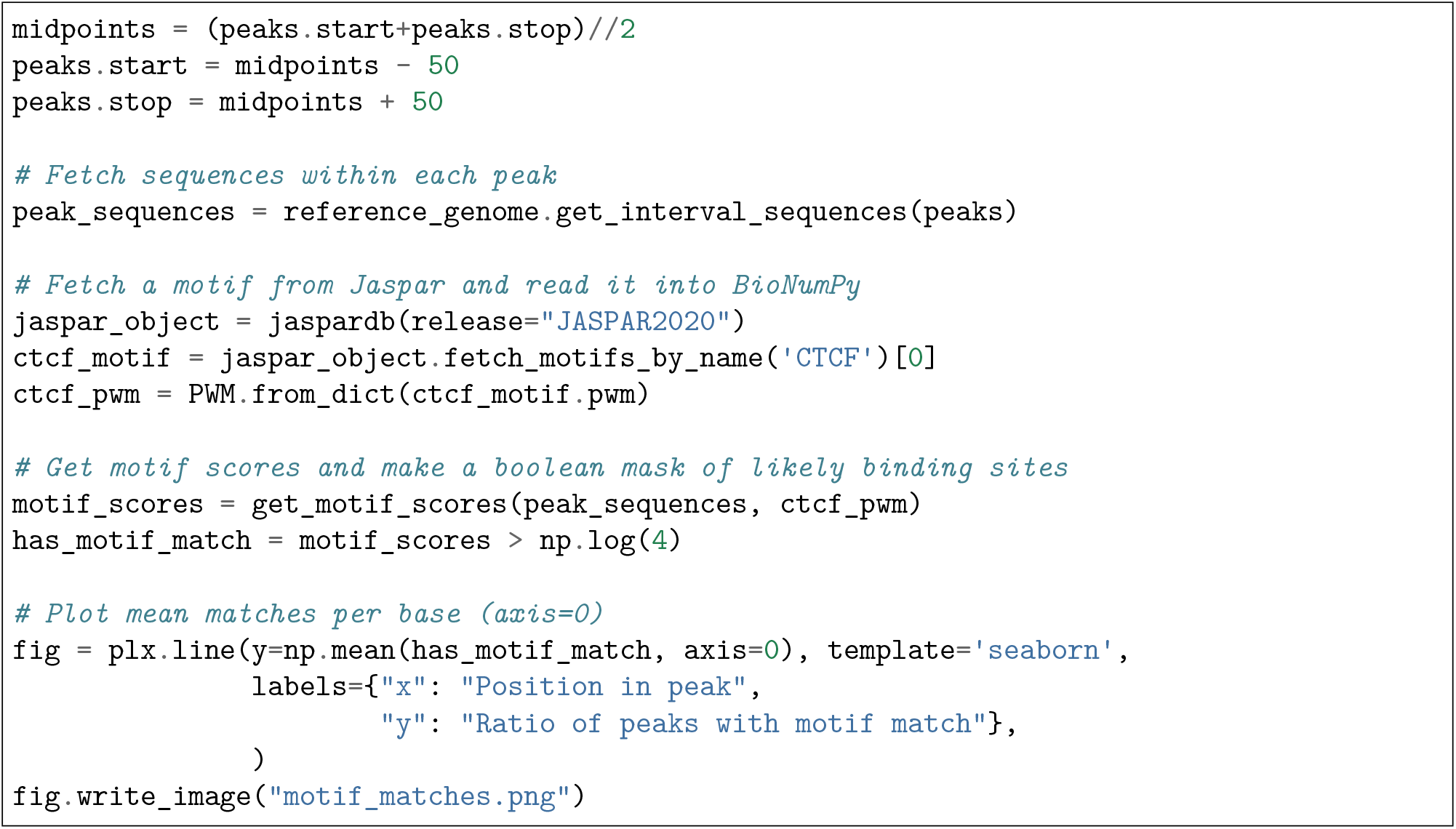

### Implementation details

#### Data representation

BioNumPy internally stores sequence data (e.g. nucleotides or amino acids) as numeric values, allowing the use of standard NumPy arrays for data representation and processing. A key way in which BioNumpy achieves high performance is by storing multiple data entries in shared NumPy arrays. To illustrate the benefit of this approach, consider the example where we want to count the number of Gs and Cs in a large set of DNA sequences. With existing Python packages like BioPython and Biotite, this must be done by iterating over the sequences using Python for-loops, which is slow when the number of sequences is large. BioNumPy, however, stores all sequences in only one or a few shared NumPy arrays (Figure 3a), meaning that vectorized NumPy operations can be used to do the counting in a fraction of the time.

Storing multiple elements in shared arrays is trivial if the elements all have the same size, since a matrix representation can be used. However, for biological data, it is common that data elements vary in size. For instance, sequences in FASTA files are rarely all of the exact same size. BioNumPy uses the RaggedArray data structure from the npstructures package (https://github.com/bionumpy/npstructures, developed in tandem with BioNumPy) to tackle this problem (Figure 2). The RaggedArray can be seen as a matrix where rows can have different lengths. The npstructures RaggedArray implementation is compatible with most common NumPy operations, like indexing (Figure 3b), vectorized operations (Figure 3c), and reductions (Figure 3d). As far as possible, objects in BioNumPy follow the array interoperability protocols defined by NumPy (https://numpy.org/doc/stable/user/basics.interoperability.html)

**Fig. 2:**
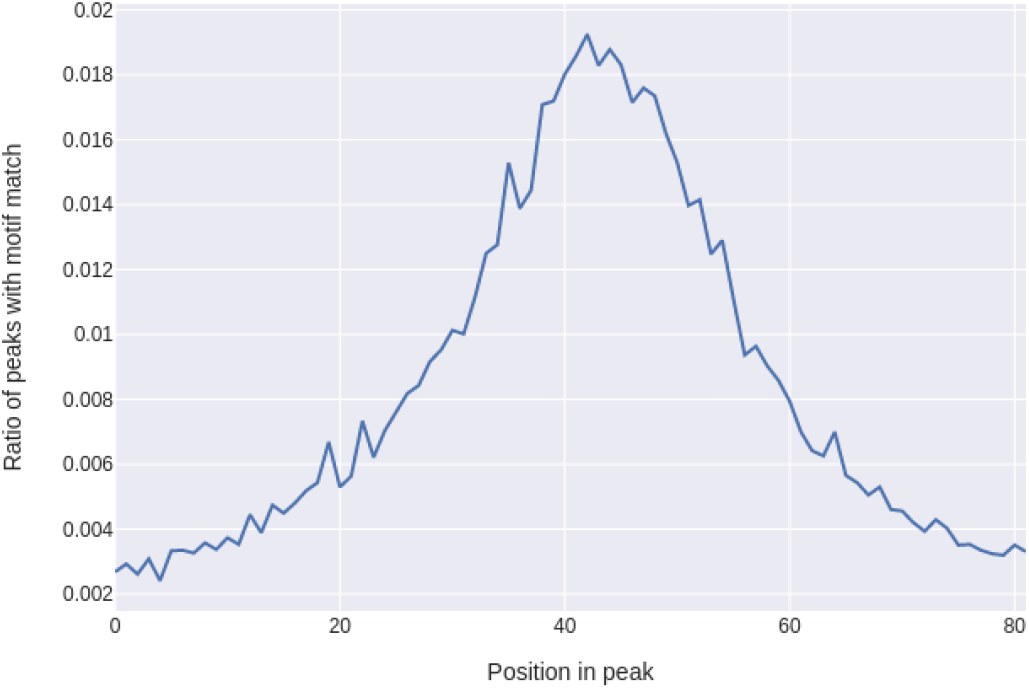
Plot generated in Example 2. Showing ratio of peaks with a motif match per base position, with an enrichment at the centre of each peak as one would expect.

**Fig. 3:**
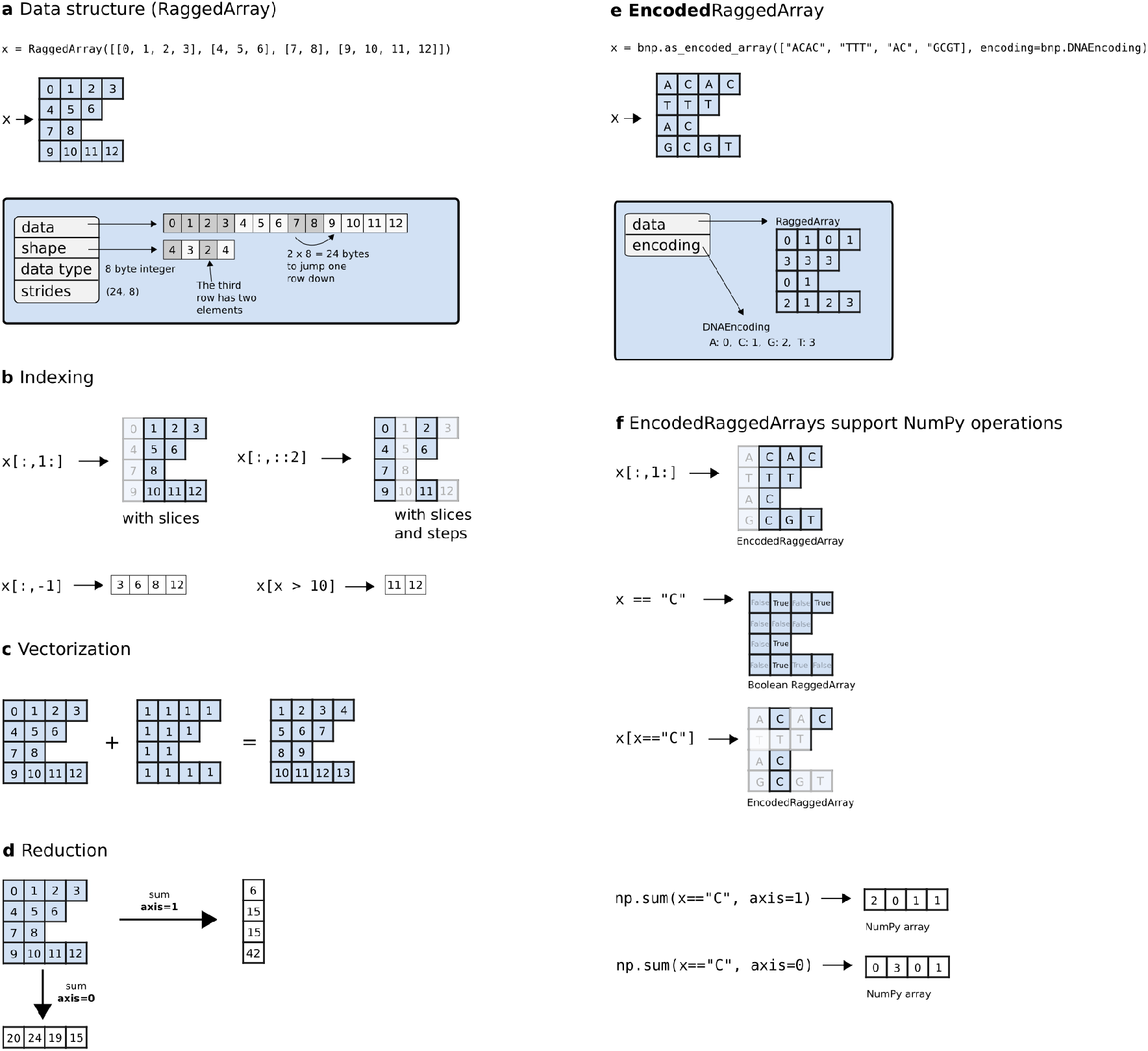
Overview of the RaggedArray and EncodedRaggedArray data structures. A RaggedArray is similar to a NumPy array/matrix but can represent a matrix consisting of rows with varying lengths (a). This makes it able to efficiently represent data with varying lengths in a shared data structure. A RaggedArray supports many of the same operations as NumPy arrays, such as indexing (b), vectorization (c) and reduction (d). An EncodedRaggedArray is a RaggedArray that supports storing and operating on non-numeric data (e.g. DNA-sequences) by encoding the data and keeping track of the encoding (e). An EncodedRaggedArray supports the same operations as RaggedArrays (f). This figure is an adopted and modified version of Figure 1 in [6] and is licensed under a Creative Commons Attribution 4.0 International License (http://creativecommons.org/licenses/by/4.0/).

#### Development

BioNumPy has been developed following the principles of continuous integration and distribution [13]. The codebase is thoroughly and automatically tested through an extensive collection of unit tests, application tests, integrations tests and property-based tests [14]. New code changes are automatically benchmarked and tested before being automatically published, ensuring that updates can be frequent, while high code quality is maintained. This makes it safe and easy to allow contributions from new contributors, which is important for longevity and community adoption of the package.

## Discussion

We have presented a new Python package, BioNumPy, for efficient representation and analysis of biological datasets. We have shown that BioNumPy is usually considerably faster than both vanilla Python scripts and commonly used Python packages for performing similar tasks. BioNumPy also has comparable efficiency to commonly used efficient tools written in C/C++.

While BioNumPy is fast on basic operations such as kmer counting and getting reverse complements of reads, we want to emphasise that BioNumPy is not specifically designed for standard tasks where tailored and highly optimised tools already exists [11]. BioNumPy is instead meant to be used as a library inside Python, and is useful when one e.g. wants to perform multiple operations on a dataset, explore or play around with datasets, or perform analyses that integrate multiple datasets in novel ways. We also invite the community to develop a broad variety of functionality for dedicated purposes with BioNumPy as an internal workhorse.

As shown in Figure 1, BioNumPy is not always faster than vanilla Python code, e.g. for the case where one is only reading a FASTA file, subsampling the sequences and writing the results back to file. The reason is that although BioNumPy reads all data into NumPy-arrays that can be efficiently subsampled, BioNumPy performs operations beyond the vanilla Python implementation, such as validating, encoding and representing the data efficiently. These additional steps come handy when you want to do more operations on the data, such as combining it with other datasets or querying it in different ways. An example of a case where BioNumPy gives considerable speedup over native tools is the problem of computing the Jaccard similarity index between all pairs of a set of bed-files. Since BioNumPy can keep all files in memory, it is considerably faster than dedicated packages like BEDTools, which needs to read each bed-file from disk every time a pair of BED-files are to be compared.

Many common bioinformatics tasks are today typically performed as a series of bash commands, using a combination of sed, AWK, Grep, Perl and/or other native unix utility tools. As an example, consider the following bash-code for converting from FASTQ to FASTA:

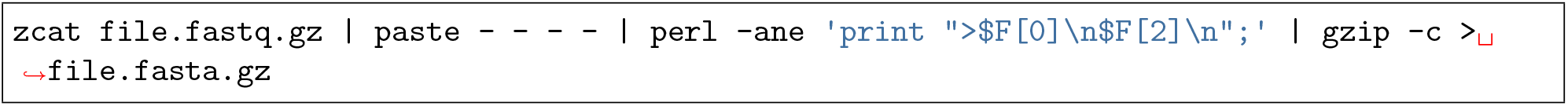

While such commands often yield fast results, there are in our opinion several drawbacks to this approach, which a package like BioNumPy addresses. First, bash commands are difficult to read and understand, which increases the chance for errors. Second, since such commands typically come in the form of ad-hoc scripts, these are usually not version controlled, not tested and thus not reproducible (each person typically has their own script). Third, it is inconvenient to write unit tests, defensive assertion code or do runtime inspection/debugging on such bash scripts, meaning that logical errors can easily go unnoticed. A better solution is thus to instead use specialised tools, such as e.g. seqtk or BioNumpy. The task of converting from FASTQ to FASTA can be done like this in BioNumPy:

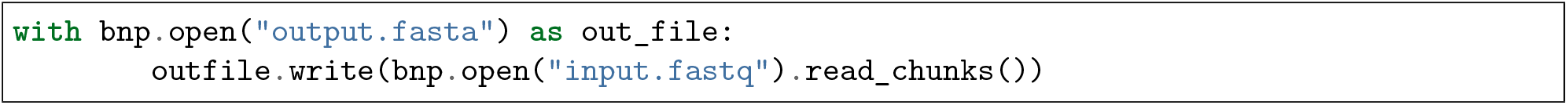

Since BioNumPy is flexible in its input, it works well with existing packages and solutions for fetching data from databases, e.g. in combination with the various BioPython modules for downloading data from databases like Encode [15] and Jaspar [12]. This ease of interoperability is also the reason why we have limited the scope of BioNumPy to not including modules for e.g. fetching data from online databases.

It can be speculated that the difficulty of writing efficient code for large-scale analyses in Python is an important reason why a lot of central bioinformatics tools are instead written in low-level and harder to learn languages like C or C++ [9, 11]. The fact that tools are written in such languages means that the large majority of bioinformaticians and computational biologists - who are typically only familiar with bash, R, and/or Python - are not able to easily contribute to the development of tools or understand/learn the internal workings of the methods they use. This limits transparency of bioinformatics research, and is also a broader problem since the continually growing size of biological data necessitates fast and efficient tools and libraries. Our hope is that BioNumPy is able to bridge this gap by making it possible for anyone to more easily work with large biological datasets in Python.

